# Small LEA proteins as an effective air-water interface protectant for fragile samples during cryo-EM grid plunge freezing

**DOI:** 10.1101/2024.02.06.579238

**Authors:** Kaitlyn M. Abe, Ci Ji Lim

**Affiliations:** Department of Biochemistry, University of Wisconsin-Madison, Madison, WI 53706

## Abstract

Sample loss due to air-water interface (AWI) interactions is a significant challenge during cryo-electron microscopy (cryo-EM) sample grid plunge freezing. We report that small Late Embryogenesis Abundant (LEA) proteins, which naturally bind to AWI, can protect samples from AWI damage during plunge freezing. This protection is demonstrated with two LEA proteins from nematodes and tardigrades, which rescued the cryo-EM structural determination outcome of two fragile multisubunit protein complexes.

## Main

The ‘resolution revolution’ of cryogenic-electron microscopy (cryo-EM) marks a significant shift in the field of structural biology^1^. However, the continued growth of cryo-EM single particle analysis (SPA) structural biology faces an unexpected problem at the step of sample grid preparation — sample damage^1–4 1,5,6^. The standard method for cryo-EM SPA sample grid preparation is plunge freezing, where small volumes of sample are applied onto a pretreated cryo-EM grid, blotted, and rapidly plunge-frozen into a cryogen such as liquid ethane. This process forms a very thin layer (∼50-100 nm) of sample-embedded vitreous ice for transmission electron microscopy imaging^7,8^. After the blotting and before the rapid freezing step, the sample exists as a thin aqueous film with a high surface-to-volume ratio. During this transitional phase (typically in seconds), the proteins in the aqueous solution collide with the air-water interface (AWI) multiple times before the sample hits the cryogen for vitrification. Adsorption to the AWI can destroy the protein’s structural integrity, causing it to denature, or disintegrate^3,5,9,10^ **(Fig. 1a)**. In a scenario where the protein maintains its structure after AWI adsorption, the sample interaction with the AWI may be biased, leading to preferred sample orientations. A non-uniform representation of the protein along the imaging axis results in an anisotropic 3D reconstruction of the sample^1,10–12^.

**Figure 1.**
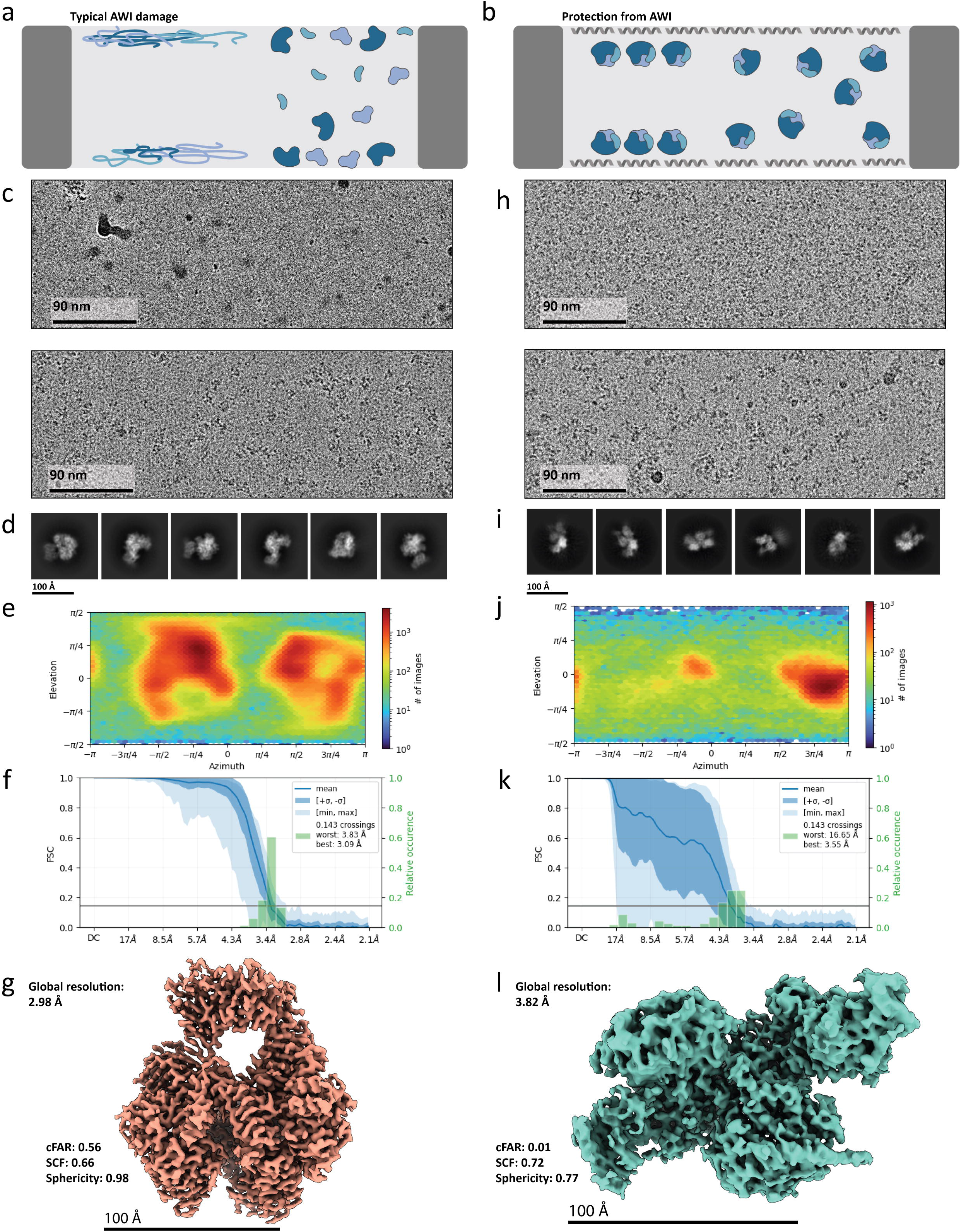
Nematode AavLEA1 effectively shields PP and PRC2 from AWI damage. **(a)** Cartoon illustration of typical outcomes when protein samples (blue) interact with the air-water interface (AWI) before vitrification. AWI damage can lead to complete denaturation (left) or partial subunit dissociation (right). **(b)** Cartoon model of the proposed LEA protein sample protection mechanism. LEA proteins form a protective protein layer at AWI, preventing larger protein samples from interacting with the AWI. LEA proteins shown as grey α-helices and protein complexes in blue. The protected protein samples may be intact but adopt biased orientations (left), or evenly distributed (right) in ice. **(c)** Representative micrographs of human DNA polymerase-α-primase (PP) with no additives (top) or with AavLEA1 at 1:40 molar ratio (bottom). **(d)** Representative two-dimensional (2D) class averages of PP with AavLEA1 (1:40). **(e)** Particle orientation distribution analysis of the PP-AavLEA1 (1:40) cryo-EM map. **(f)** Fourier-shell correlation (FSC) curve of the PP-AavLEA1 (1:40) cryo-EM map. **(g)** Cryo-EM map of PP-AavLEA1 (1:40) with a global resolution of 3.0 Å (reported at 0.143 FSC). **(h)** Representative micrographs of human Polycomb Repressive Complex 2 (PRC2) without (top) or with AavLEA1 added at 1:40 molar ratio (bottom). **(i)** Representative 2D class averages of PRC2 with AavLEA1 (1:40). **(j)** Particle orientation distribution analysis of the PRC2-AavLEA1 (1:40) cryo-EM map. **(k)** FSC curve of the PRC2-AavLEA1 (1:40) cryo-EM map. **(l)** Cryo-EM map of PRC2-AavLEA1 (1:40) dataset with a global resolution of 3.8 Å.

To mitigate the AWI problem, researchers have developed a variety of solutions. These include the addition of mild detergents to the sample^6^, the use of chemical crosslinking^3,13,14^, and adsorbing or tethering samples to surfaces to prevent sample contact with the AWI^12,15–24^. More sophisticated advanced techniques involve rapidly freezing the sample before the proteins can diffuse to the AWIs^25–27^. However, these methods are often sample-specific, technically challenging, or cost-prohibitive. In this study, we report a simple, effective, and economical approach for cryo-EM SPA users to circumvent issues related to AWI sample damage. In this work, we show that late embryogenesis abundant (LEA) proteins, which naturally bind to the AWI^28^, can be utilized to shield samples from AWI-induced damage during cryo-EM grid plunge freezing (**Fig. 1a**). This new approach enables us to determine cryo-EM SPA structures of challenging multisubunit protein complexes at higher or comparable resolution than those previously reported using complicated AWI damage deterrent methods.

The AWI protective function of LEA proteins is demonstrated with proteins from two organisms: AavLEA1^28–31^ from the nematode *Aphelenchus avenae* and RvLEAM^20^ from the tardigrade *Ramazzottius varieornatus*, also known as water bears. These proteins are small — approximately 16 kilodaltons (kDa) for AavLEA1 and 30 kDa for RvLEAM. Hence, we deduced that their small size enables rapid diffusion in solution, allowing them to reach the AWI faster than the larger protein samples of interest (typically, they are several hundred kDa in size). The LEA proteins will be stably adsorbed to the AWI due to their natural AWI affinity, thus forming a protective protein layer preventing the samples from contacting the AWI (**Fig. 1a**). In this study, RvLEAM was truncated (hereafter termed RvLEAM1_short_) to approximately 15 kDa to further enhance diffusion efficiency (refer to the methods section for details).

To evaluate the AWI protective effects of the LEA proteins, we used two protein complexes that are AWI-sensitive as model systems: human DNA polymerase-α-primase (PP)^32–34^ and Polycomb repressive complex-2 (PRC2)^35,36^. It is important to note that in this study, both complexes were tested in their apo-state, and all data collection was performed using a 200 keV electron microscope with a direct electron detector. Reported cryo-EM structures of apo-state PP complexes were solved using chemically crosslinked samples^34^, suggesting that the PP particles fell apart due to AWI damage during plunge freezing. Indeed, in our hands, we saw no discernable PP particles using standard plunge freezing conditions with a Vitrobot (Mark IV, Thermo Fisher Scientific, USA) (**Fig. 1c**). Another arguably more challenging structure to solve for cryo-EM SPA is that of PRC2. Reported PRC2 cryo-EM structures were solved by highly specialized strategies using either chemically crosslinked samples on carbon-support grids^37,38^ or biotinylated samples tethered onto Streptavidin-crystal grids^13,39–42^.

We found that the addition of AavLEA1 or RvLEAM1_short_ to PP and PRC2 prior to freezing eliminate the need for the abovementioned complicated methods, allowing us to solve their structures at comparable or higher cryo-EM map resolutions (**Fig. 1c-l**). The optimal range of addition is found to be between 1:6-1:40 molar ratio of the sample to the LEA proteins (**Extended Data Fig. 1**). For example, the addition of AavLEA1 at a 1:40 PP:AavLEA1 molar ratio before plunge freezing led to a much more positive outcome than the PP sample grid without any additives; we saw discernable and homogeneously sized particles (**Fig. 1c**). Multiple high-resolution two-dimensional (2D) class averages of PP were obtained from these particles (**Fig. 1d**). Subsequent image processing led to a 3.0 Å global resolution cryo-EM map of the apo-state PP (**Fig. 1e-g, and Extended Data Fig. 2**). This cryo-EM map is the highest resolution reported for the human apo-state PP; the previous X-ray structure was solved at 3.6 Å^43^ and a chemically crosslinked cryo-EM structured determined at a 3.8 Å global resolution^34^. Most importantly, we demonstrated that LEA proteins, AavLEA1 in this case, can be used to “resuscitate” an unviable cryo-EM project and achieve high-resolution cryo-EM structures.

Initial concerns with the AavLEA1 strategy are 1) elevated background signal in the micrograph from the AavLEA1 addition and 2) AavLEA1 distorting the conformation of the sample of interest. In our experiments, we found no significant increase in the background signal (**Fig. 1c**), even at the highest tested AavLEA1:sample ratio of 40:1. If there were any, it did not affect image alignment or hindered high-resolution cryo-EM structure determination, as can be seen from our high-resolution map of PP (**Fig. 1g**). In addition, we did not find any extra map density that would indicate AavLEA1 binding to PP nor a significant change in the conformation (R.M.S.D of 1.1 Å when aligned to 5EXR^43^, see **Extended Data Fig. 3**). In short, AavLEA1 addition did not alter PP apo-state conformation.

CHAPSO, a zwitterionic detergent, can protect protein samples from AWI damage during plunge freezing^44,45^. In our hands, it has been an effective strategy for determining the high-resolution cryo-EM structures of PP-related complexes^33,46^. Hence, we want to compare AavLEA1’s efficiency in obtaining a high-resolution cryo-EM structure against CHAPSO. To this end we determined a 3.5 Å cryo-EM map of apo-state PP using CHAPSO (**Extended Data Fig. 4**). By visual inspection, we found no discernable differences between the two cryo-EM maps. Their particle image quality is also approximately similar, as indicated by ResLog analysis^47^ (**Extended Fig. 5**). Thus, the particle quality in the AavLEA1 dataset is comparable to that of the CHAPSO dataset.

However, the use of CHAPSO or similar detergents requires a significantly higher concentration of samples^48^ — over 4 mg/mL or approximately 13 μM for a 300 kDa protein sample. This presents a challenge for precious samples. In contrast, with AavLEA1, we only use around 1-1.5 μM of sample protein, which is more than ten times less than what is required for detergents. Therefore, AavLEA1 offers a valuable option when sample is limiting. In our hands, sample aggregation is a common problem from adding detergents. Another benefit of AavLEA1 is the early detection of potential interaction with the protein sample. If interaction occurs, it is more likely to be detected in a preliminary low-resolution cryo-EM map given AavLEA1’s size of ∼15 kDa. This is unlike CHAPSO, a small molecule that requires sub-4 Å resolution maps for detection^45^.

We saw similar results for PRC2 with the addition of AavLEA1 (1:40 PRC2:AavLEA1 molar ratio) — Adding AavLEA1 resulted in discernable and homogeneously sized particles (**Fig. 1h**). Without additives, we saw smaller sized particles in the micrographs, indicating broken PRC2 complexes which is consistent as previously reported^13^. Subsequent cryo-EM image processing led to a 3.62 Å global resolution cryo-EM map of PRC2 (**Fig. 1i-l**). To our knowledge, this is the first high-resolution cryo-EM map of PRC2 that was obtained without the use of chemical crosslinking or adsorption to a carbon surface. This achievement further exemplifies the efficacy of AavLEA1 in facilitating the determination of high-resolution cryo-EM structures of protein samples sensitive to the air-water interface.

Notably, the AavLEA1-derived PRC2 cryo-EM map suffered from preferred particle orientation (**Fig. 1j & k**). In retrospect, the AavLEA1-derived PP cryo-EM map also has a certain degree of biased particle orientations, although this did not prevent us from obtaining an isotropic map (**compare Fig. 1g, and Extended Data Fig. 4**). If the LEA proteins are indeed providing protection to the protein samples from AWI interactions, one would expect the sample particles to exhibit a random yet evenly distributed orientation, like the one observed with the PP cryo-EM map in the presence of CHAPSO (**Extended Data Fig. 4c**). A possible explanation to why there is a biased but different particle orientation in the AavLEA1-PP and -PRC2 datasets is that the AavLEA1 protective layer at the AWI interacts with the protein samples in a manner specific to each sample. Nonetheless, these interactions do not appear to compromise the integrity of both samples, as evidenced by their high-resolution cryo-EM maps. In addition, problems associated with preferred particle orientation can be mitigated by employing tilted-stage data collection methods^11^.

After establishing AavLEA1’s efficacy in AWI protection, we investigated whether the tardigrade LEA protein, RvLEAM, exhibits similar protective qualities. Indeed, RvLEAM_short_ similarly protects PP (**Extended Data Fig. 6**) and PRC2 (**Fig. 2a and Extended Data Fig. 7**) from AWI damage during plunge-freezing, suggesting that AWI protection is a universal characteristic of LEA proteins. We also noticed that the cryo-EM maps of PP and PRC2 obtained from the RvLEAM_short_ datasets had particle orientation distributions comparable to those seen in the AavLEA1 datasets (**refer to Fig. 1j and 2a**). This similarity in particle orientation distribution between the two datasets suggests a shared AWI protection mechanism by the LEA proteins. Both AavLEA1 and RvLEAM_short_ are predicted to form amphiphilic alpha helical structures, with the hydrophobic face oriented towards the air at the AWI and the hydrophilic side (negatively charged) facing the aqueous solution^49,50^. This offers a simple explanation for the observed similarity in particle orientation distributions when samples are exposed to either of the LEA proteins — both LEA proteins present a negatively-charged surface (water-facing) at the AWI.

**Figure 2.**
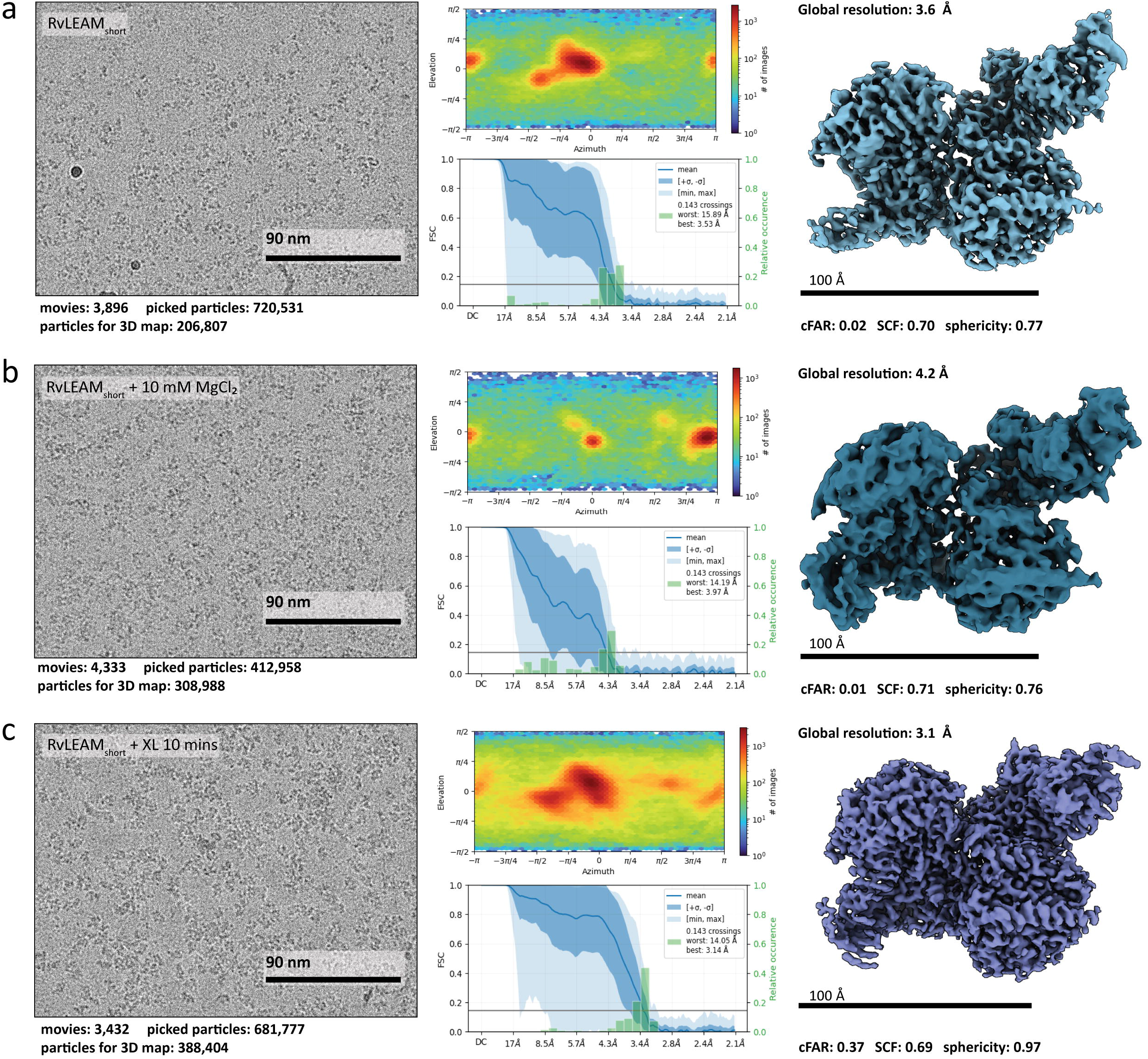
Tardigrade RvLEAM_short_ protects PRC2 from AWI damage and can be used in conjunction with other buffer additives to diversify particle orientation distribution. **(a)** PRC2-RvLEAM_short_ (1:6 molar ratio) cryo-EM single-particle analysis: representative micrograph, particle orientation analysis, FSC curve, and the cryo-EM map with a 3.6 Å global resolution. **(b)** PRC2-RvLEAM_short_ (1:6 molar ratio) with 10 mM MgCl_2_ cryo-EM single-particle analysis: representative micrograph, particle orientation analysis, FSC curve, and the cryo-EM map with a 4.2 Å global resolution. **(c)** PRC2-RvLEAM_short_ (1:6 molar ratio) with 10 minutes of glutaraldehyde crosslinking cryo-EM single-particle analysis: representative micrograph, particle orientation analysis, FSC curve, and the cryo-EM map with a 3.1 Å global resolution. All global resolution values are reported at 0.143 FSC threshold.

Next, we explored conditions to improve PRC2 particle orientation distribution in the presence of RvLEAM_short_ (**Fig. 2b-d**). First, we added divalent cation, MgCl_2_, to attempt to neutralize the negatively charged surface presented by RvLEAM_short_ at the AWI, with the hope that this strategy will change the PRC2 particle orientation distribution. Indeed, we observed a significant shift in the PRC2 particle orientation distribution, but the distribution is still heavily biased (**Fig. 2b**). Next, we crosslinked the PRC2 samples with glutaraldehyde before freezing it with RvLEAM_short_ (**Extended Data Fig. 8**). The carbonyl ends of glutaraldehyde form Schiff bases with the positively charged groups on the protein surface. This can neutralize the positively charged surface of PRC2 and change a sample’s particle orientation distribution^51^. Indeed, we found additional new particle views in the crosslinked PRC2-RvLEAM_short_ dataset and obtained a 3.10 Å cryo-EM map of PRC2 with improved isotropy (**Fig. 2c**). A milder crosslink treatment yielded a similar outcome (**Extended Data Fig. 9**). We also noticed significantly more intact PRC2 particles during image processing for these crosslinked datasets. The above suggests cryo-EM users can combine chemical crosslinking and LEA protein AWI protection for challenging cryo-EM samples.

## Discussion

In summary, we showed that LEA proteins is an effective sample protectant for AWI damage in cryo-EM grid preparation. By simply adding AavLEA1 or RvLEAM_short_ to samples before plunge freezing, we were able to determine the cryo-EM structures of both PP and PRC2 at comparable, or better resolution than those previously reported using more challenging and complicated anti-AWI damage strategies^13,34,37–42^. The LEA protein additives provide a simple, robust, and cost-effective anti-AWI damage solution that any cryo-EM labs and facilities in the world can readily incorporate into their existing workflow.

## Methods

### Expression and purification of AavLEA1 and RvLEAM_short_

The expression plasmid for HIS-tagged AavLEA1 was obtained from the Addgene plasmid repository [pET15b-AavLEA1 was a gift from Claude Férec (Addgene plasmid # 53093)]^52^ The expression plasmid for HIS-tagged RvLEAM_short_ was made by inserting a truncated cDNA from pEThT-RvLEAM^49^ [pEThT-RvLEAM was a gift from Takekazu Kunieda (Addgene plasmid # 90033)], into the pET15b vector. RvLEAM_short_ encodes residues 58-181 of RvLEAM (A0A0E4AVP3.1). Recombinant AavLEA1 and RvLEAM_short_ were expressed in *Escherichia coli* BL21 (DE3) cells. Briefly, a single bacterial colony with the transformed plasmid was inoculated in 2 mL Luria Bertani broth (LB) with 100 μg mL^-1^ carbenicillin and grown overnight at 37°C. This starter culture was then used to inoculate 1 L of LB supplemented with antibiotics. Once the culture reached an absorbance (A_600_) of 0.6, gene expression was induced with 0.1 mM isopropyl-β-D-thiogalactopyranoside (IPTG) for 16 hours at 12°C, shaking at 230 rpm.

Next morning, the cells were harvested by centrifugation and resuspended in lysis buffer (50 mM HEPES pH 7.5, 300 mM NaCl, 10 mM imidazole, 1 mM DTT/TCEP, 1 mM PMSF). The cells were lyzed by sonication and then centrifuged to remove debris. Pre-equilibrated nickel-NTA resin (Qiagen, 50 mg/mL binding capacity) was added to the clarified lysate and the lysate-resin slurry stirred in a beaker with a stir bar for 1 hour at 4°C. The protein-bound resin was washed three times, each time using 50 mL of lysis buffer. After the final wash, 10 mL of elution buffer (wash buffer supplemented with final 250 mM imidazole) was added to the resin, and the eluted protein collected using a gravity flow column. The proteins were concentrated to ∼500 uL using a 3 kDa MWCO spin column before it is injected into a Superdex 75 10/300 size-exclusion chromatography (SEC) column (Cytiva, USA) that is pre-equilibrated with SEC buffer (50 mM HEPES pH 7.5, 300 mM NaCl, 1 mM DTT/TCEP, 10% glycerol). Eluted fractions were collected and analyzed by SDS-PAGE. Fractions containing the proteins were pooled, concentrated and then snap-frozen as 5-10 μL aliquots. The frozen aliquots are stored in a -80°C freeze until usage.

### Production of recombinant human DNA Polymerase alpha-primase and Polycomb repressive complex 2

Recombinant human DNA Polymerase alpha-primase were expressed in insect cells using baculovirus infection and purified as previously reported^46^. Purified recombinant human PRC2 protein complexes were generous gifts from Dr. Thomas Cech at the University of Colorado Boulder. The expression and purification protocols of PRC2 has be described previously^40^.

### PRC2 glutaraldehyde crosslinking

Approximately 2 μM PRC2 were incubated with a 0.1% (v/v) final glutaraldehyde concentration for 2- and 10-minute time intervals. After each interval, an aliquot is removed and quenched with 80 mM Tris-HCl to stop the reaction. SDS-PAGE was used to determine the crosslinking efficiency of the PRC2 samples at various incubation time point.

### Cryo-EM sample grid preparation and plunge-freezing

All samples used were thawed just prior to cryo-EM grid preparation. Holey carbon cryo-EM grids, either Quantifoil R 1.2/1.3 300 mesh Au or C-flat R 1.2/1.3 300 mesh Au were glow discharged on a PELCO EasiGlow glow-discharge unit (15 mA for 30 seconds with 10 second hold). The protein samples were diluted to working concentration right before grid application in 25 mM HEPES pH 7.5, 150 mM NaCl, 1 mM TCEP. Where indicated, buffer was supplemented with CHAPSO or MgCl_2_. Then 3.5 μL of the sample was applied to the glow discharged grid. The grid was then blotted for 4 to 6 seconds at 4°C and 95% humidity, and then plunge frozen into liquid ethane using a Vitrobot Mark IV (Thermo Fisher, USA).

For all conditions except those specified, 1 – 1.5 μM of PP or PRC2 were used with the indicated molar ratio of AavLEA1 or RvLEAM_short_. For conditions using PP and 4 mM CHAPSO, 13.8 μM PP was used.

### Cryo-EM data collection

All data collections and screening were done on a Talos Arctica 200 kV TEM (Thermo Fisher, USA) with a Gatan BioQuantum K3 direct electron detector (Gatan, USA). Data screening and acquisition software used was either EPU (Thermo Fisher, USA) or SerialEM^53^. All cryo-EM datasets were collected at a pixel size of 1.064 Å/pixel with a total dose of 50 e^-^Å^-2^ across 40 frames. CDS counting mode was used with energy filter inserted at 20 eV. Defocus range used was -1 to -2.5 μm.

### Cryo-EM data processing

For all datasets, image processing was performed using cryoSPARC^54^. Briefly, movies were motion corrected and the ensuing micrographs had their contrast transfer function (CTF) estimated. The CTF values were used to curate a set of micrographs that are suitable for high-resolution single-particle analysis. Details of subsequent image processing steps for each dataset are as follows:

#### 1.5 μM PP, 12 μM AavLEA1 (1:8) dataset

A total of 2,764 movies were collected. After micrograph curation 2,112 movies remained and 1,724,984 particles were extracted at 4x binning (4.3 Å per pixel). After 2D classification, 785,653 particles proceeded to ab-initio reconstruction and sorted into four separated reference-free 3D classes. Particles underwent another round of ab-initio modeling and separated into two classes. The intact complex was sorted into one of the two classes (239,029 particles, 75%) and non-uniform refinement of this class with per-particle CTF refinement resulted in a global resolution (reported at Fourier shell correlation of 0.143) of 3.6 Å.

#### 1.5 μM PP, 60 μM AavLEA1 (1:40) dataset

A total of 4,403 movies were collected. After micrograph curation, 500 movies were initially used with a total of 340,430 particles extracted at 4x binning. From 2D classification, 201,337 particles were selected and re-extracted at original pixel size. Particles then proceeded to ab-initio reconstruction and were sorted into four separated reference-free 3D classes. Two of the four classes resulted in intact particles, which were verified through non-uniform refinement of the combined classes using 165,472 particles, resulting in a 3.8 Å structure. Particles were extracted from the remaining 3,771 movies and binned 4x, resulting in 2,668,602 particles. These particles were then sorted into 2D classes, and the selected 1,438,074 particles underwent ab-initio modeling. Selected particles underwent another round of ab-initio modeling with two classes. One of the two classes showed intact particles, and those 1,009,026 particles were sent to non-uniform refinement yielding a 3.43 Å global resolution. CryoSparc global and local CTF refinement jobs were run followed by further filtering, resulting in 988,417 particles. These particles were then extracted at the original pixel size. All resulting particles underwent non-uniform refinement, reference motion correction, and heterogeneous refinement. 856,205 particles were used for a final non-uniform refinement, with a final global resolution of 3.0 Å.

The published apo-state PP (PDB: 5EXR)^43^ model was used as the initial model for real space refinement against the cryo-EM map using the Phenix software^55^. Structural alignment between the 5EXR model and the refined model was performed using the matchmaker module in the ChimeraX software^56^. The refinement statistics and validation report are available in Supplementary Table 1.

#### 1 μM PP, 6 μM RvLEAM_short_ (1:6) dataset

A total of 1,308 movies were collected. Following micrograph curation, 335,907 particles were extracted from 1,076 micrographs and binned 4x. 125,231 particles were extracted with 2D classification. These particles then proceeded to ab-initio reconstruction and split into 3D classes. The selected 74,198 particles were re-extracted at the original pixel size from 1,071 micrographs. Particles underwent another round of 2D classification and Ab-Initio 3D reconstruction. Selected particles underwent non-uniform refinement and had a final global resolution of 4.5 Å. The final resolution of this dataset is lower than our other PP datasets which is likely because this data collection contains only 1,308 movies while other PP datasets had 2,700 or more movies.

#### 13.8 μM PP, 4 mM CHAPSO dataset

A total of 4,567 movies were collected. Following micrograph curation, 1,832,455 particles were extracted from 4,510 micrographs and binned 4x. Particles underwent 2D classification where and ab-initio reconstruction. Selected particles were used to re-extract at the original pixel size (744,824 particles). Extracted particles were then sorted into 2D classes underwent non-uniform refinement with per-particle CTF refinement and reference motion correction, with a final global resolution of 3.4 Å with 674,793 particles.

#### 1.5 μM PRC2, 60 μM AavLEA1 (1:40) dataset

A total of 2,843 movies were collected. Following micrograph curation, 687,121 particles were extracted from 1,642 micrographs and binned 4x. Particles underwent 2D classification and ab-initio reconstruction. Selected classes were re-extracted at original pixel size, yielding 150,359 particles. The re-extracted particles underwent non-uniform refinement with per-particle CTF refinement and resulted in a global resolution of 3.8 Å.

#### 1 μM PRC2, 12 μM AavLEA1 (1:12) dataset

A total for 4,356 movies were collected. Following micrograph curation, 2,073,578 particles were extracted from 4,112 movies and binned 4x. Particles were sorted into 2D classes followed by ab-Initio 3D classes. One of the four classes showed an intact complex, so the 275,416 particles from this class were re-extracted at the original pixel size from 4,110 movies. Re-extracted particles underwent another round of ab-initio 3D reconstruction. The final 238,045 particles underwent non-uniform refinement, with a final global resolution of 3.7 Å.

#### 1 μM PRC2, 6.7 μM RvLEAM_short_ (∼1:6) dataset

A total of 3,896 movies were collected. Following micrograph curation, 1,766,944 particles were extracted from 3,637 micrographs and binned 4x. Particles were sorted into 2D classes followed by 3D classes via ab-Initio reconstruction. The selected 547,610 particles were re-extracted and subjected to ab-initio 3D reconstruction. The final 206,807 particles underwent non-uniform refinement with per-particle CTF refinement, resulting in a global resolution of 3.8 Å.

#### 1 μM PRC2, 6.7 μM RvLEAM_short_ (∼1:6) 10 mM MgCl_2_ dataset

A total of 4,333 movies were collected. Following micrograph curation, 1,755,138 particles were extracted from 3,759 micrographs and binned 4x. Particles underwent two rounds of 2D classification From here, 408,373 particles were selected and proceeded to ab-initio 3D reconstruction. The resulting 406,148 particles were re-extracted from 3,746 movies at the original pixel size and underwent another round of ab-initio reconstruction. The final 102,181 particles proceeded to non-uniform refinement with a final global resolution was 4.2 Å.

#### 1 μM PRC2 cross-linked 2 minutes, 6.7 μM RvLEAM_short_ (∼1:6) dataset

A total of 2,981 movies were collected. Following micrograph curation, 982,264 particles were extracted from 2,189 movies and binned 4x. Particles were sorted into 2D classes followed by 3D classes via ab-initio reconstruction. Selected particles were re-extracted at the original pixel size and underwent ab-initio reconstruction again. The resulting 654,585 particles were subjected to non-uniform refinement with per-particle CTF refinement, resulting in a global resolution of 3.5 Å.

#### 1 μM PRC2 cross-linked 10 minutes, 6.7 μM RvLEAM_short_ (∼1:6) dataset

A total of 3,432 movies were collected. Following micrograph curation, 2,084,816 particles were extracted from 3,295 movies and binned 4x. Particles underwent 2D classification followed by ab-initio 3D reconstruction. Selected particles were re-extracted at the original pixel size. Particles underwent non-uniform refinement, resulting in a global resolution of 3.3 Å. Global CTF refinement, reference-based motion correction and heterogeneous refinement jobs were run. 366,459 particles were used in this final round of non-uniform refinement, resulting in a global resolution of 3.1 Å.

## Data availability

The described cryo-EM maps and coordinate files have been deposited in the Electron Microscopy Data Bank and the Protein Data Bank (PDB) under the accession codes: EMD-43619 (PP - AavLEA1, 1:8 molar ratio), PDB-ID 8VY3 with EMD-43628 (PP - AavLEA1, 1:40 molar ratio), EMD-43626 (PP – RvLEAM_short_, 1:6 molar ratio), EMD-43627 (PP – CHAPSO 4 mM), EMD-43620 (PRC2 – AavELA1, 1:40 molar ratio), EMD-43621 (PRC2 – RvLEAM_short_, 1:6 molar ratio), EMD-43622 (PRC2 – RvLEAM_short_, 1:6 molar ratio, 10 mM MgCl_2_), EMD-43623 (PRC2 – AavLEA1, 1:12 molar ratio), EMD-43623 (PRC2 – RvLEAM_short_, 1:6 molar ratio, 2 min cross-link), EMD-43625 (PRC2 – RvLEAM_short_, 1:6 molar ratio, 10 min cross-link).

## Competing interests

A provisional patent has been filed for this technology with the Wisconsin Alumni Research Foundation (WARF).

## Author contributions

C.L conceived the study. K.M.A. made the recombinant proteins, cryo-EM grids, performed data collection and image processing with C.L. in support. K.M.A performed the crosslinking experiments and analysis. K.M.A. and C.L. wrote the manuscript, with the figures made by K.M.A.

## Acknowledgments

We thank Thomas Cech, Anne Gooding, and Jiarui Song at the University of Colorado Boulder for their generous gift of purified recombinant human PRC2. We also thank Qixiang He, Xiuhua Lin, and the rest of the Lim laboratory for their helpful suggestion and generous support of reagents. We thank Timothy Grant and Elizabeth Wright for their helpful suggestions. Some of this work was performed in the Cryo-EM Research Center (CEMRC) in the Department of Biochemistry at the University of Wisconsin-Madison. We thank the staff at CEMRC for their support and assistance in cryo-EM data collection. Support for this research was provided to C.L. by the National Institutes of Health (NIH), the National Institute of General Medical Sciences (R00GM131023 and DP2GM150023) and the University of Wisconsin–Madison, Office of the Vice-Chancellor for Research and Graduate Education with funding from the Wisconsin Alumni Research Foundation and the Department of Biochemistry. In addition, K.M.A is supported by a NIH T32 predoctoral fellowship (T32GM130550).

## Notes

### Competing Interest Statement

The authors have declared no competing interest.

### Summary of Updates

Methods are updated to include details of the RvLEAMshort truncation and the production of the recombinant human DNA polymerase alpha-primase and polycomb repressive complex 2.

