## Supplementary Information for "Small LEA proteins as an effective air-water interface protectant for fragile samples during cryo-EM grid plunge freezing"

**Affiliations:**

### Table of contents:

**Extended Data Figure 1.** Representative micrograph comparison of different ratios of PP and AavLEA1.

**Extended Data Figure 2.** Data processing pipeline for PP and AavLEA1 (1:40) dataset.

**Extended Data Figure 3.** Map to model fitting for PP and AavLEA1 (1:40).

**Extended Data Figure 4.** Data processing pipeline for PP and 4 mM CHAPSO dataset.

**Extended Data Figure 5.** Reslog comparison of PP and CHAPSO (4 mM) with PP and AavLEA1 (1:40)**.**

**Extended Data Figure 6.** Data processing pipeline for PP and RvLEAMshort (1:6) dataset**.**

**Extended Data Figure 7.** Data processing pipeline for PRC2 and RvLEAMshort (1:6) dataset.

**Extended Data Figure 8.** PRC2 cross-linking validation SDS-PAGE**.**

**Extended Data Figure 9.** Data processing pipeline for PRC2 and RvLEAMshort (1:6) dataset with mild cross-linking (2 minutes)**.**

**Table 1: Cryo-EM data collection, refinement, and validation statistics**

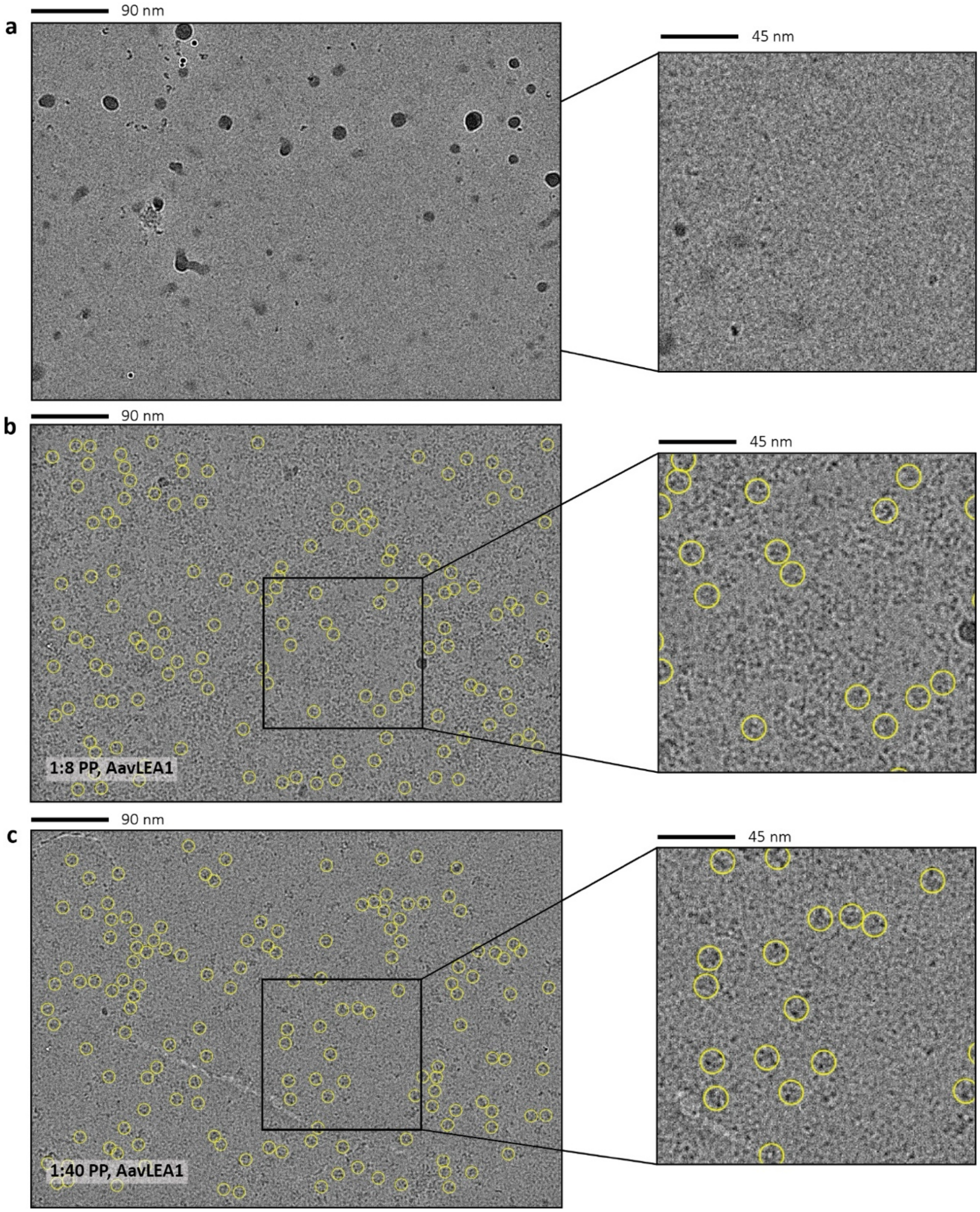

**Extended Data Figure 1 – Representative micrograph comparison of different ratios of PP and AavLEA1** Micrograph on left with zoomed-in section on right. **(a)** 1 µM PP with no added AavLEA1. No discernable intact particles of PP. **(b)** 1.5 µM PP with 12 µM AavLEA1 (1:8 molar ratio). Several particles visible that are the expected size for intact PP. **(c)** 1.5 µM PP with 60 µM AavLEA1 (1:40 molar ratio). Several particles visible that are expected size for intact PP.

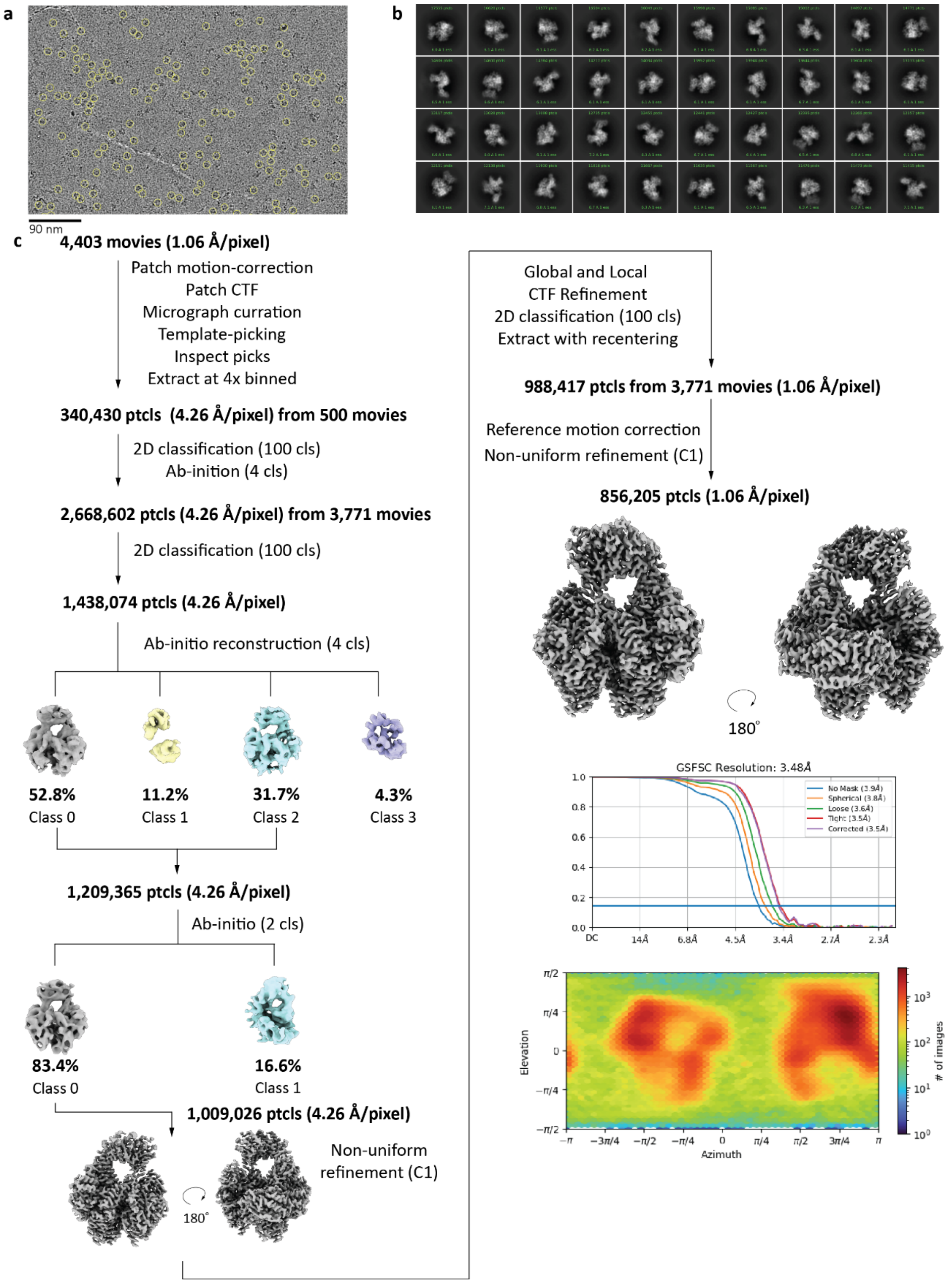

**Extended Data Figure 2 – Data processing pipeline for PP and AavLEA1 (1:40) dataset** 1.5 µM PP with 60 µM AavLEA1 **(a)** Representative micrograph. **(b)** Top 40 classes from 2D classification of PP. **(c)** Cryo-EM processing pipeline used to obtain final cryo-EM map for PP.

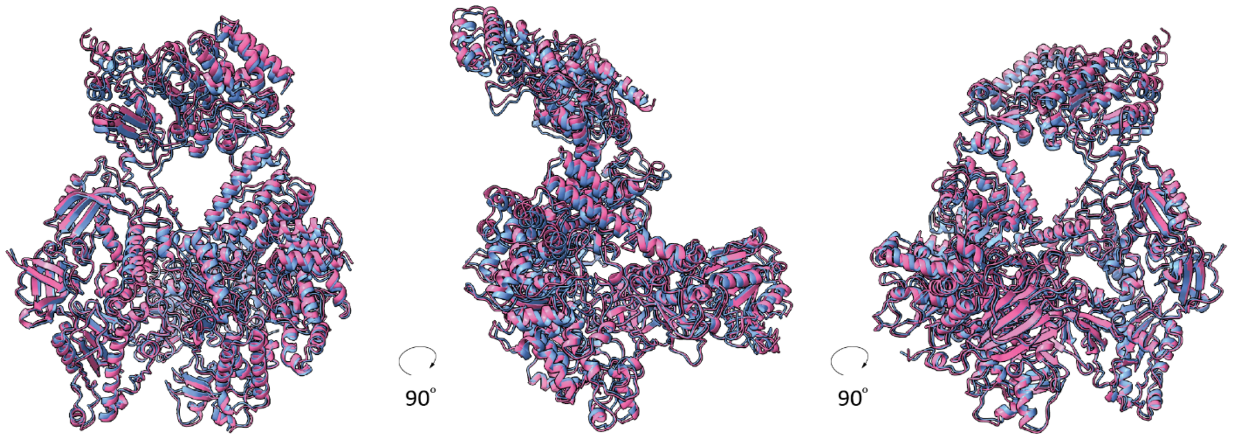

**Extended Data Figure 3 – Map to model fitting for PP and AavLEA1 (1:40)** PP model 5EXR fit into PP map from PP and AavLEA1 (1:40) dataset using Phenix. The 5EXR model is in blue and the model from the present dataset is in pink.

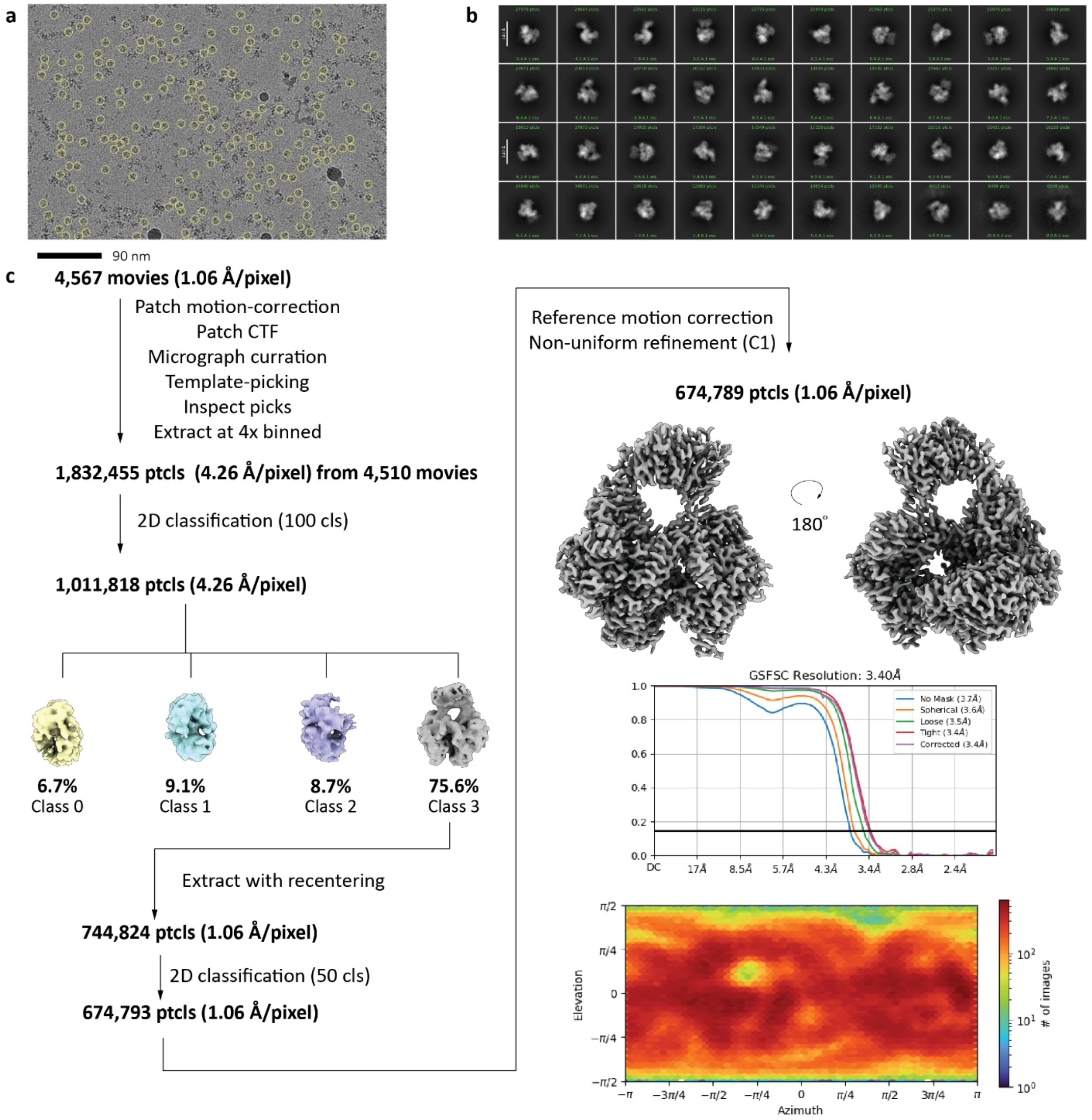

**Extended Data Figure 4 – Data processing pipeline for PP and 4 mM CHAPSO dataset** 13.8 µM PP with 4 mM CHAPSO **(a)** Representative micrograph. **(b)** Top 40 classes from 2D classification of PP. **(c)** Cryo-EM processing pipeline used to obtain final cryo-EM map for PP.

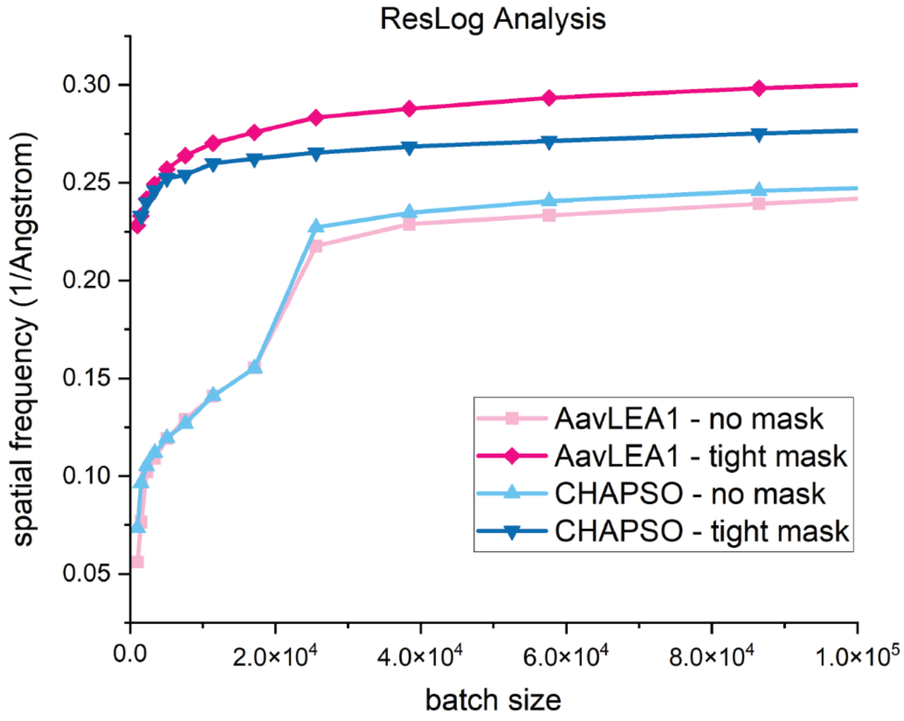

**Extended Data Figure 5 – Reslog comparison of PP and CHAPSO (4 mM) with PP and AavLEA1 (1:40)** Reslog analysis done through CryoSPARC using Fourier shell correction at 0.143. The x-axis shows the number of particles, or batch size, used in each reconstruction. The y-axis is the inverse of the resolution calculated from each batch of particles.

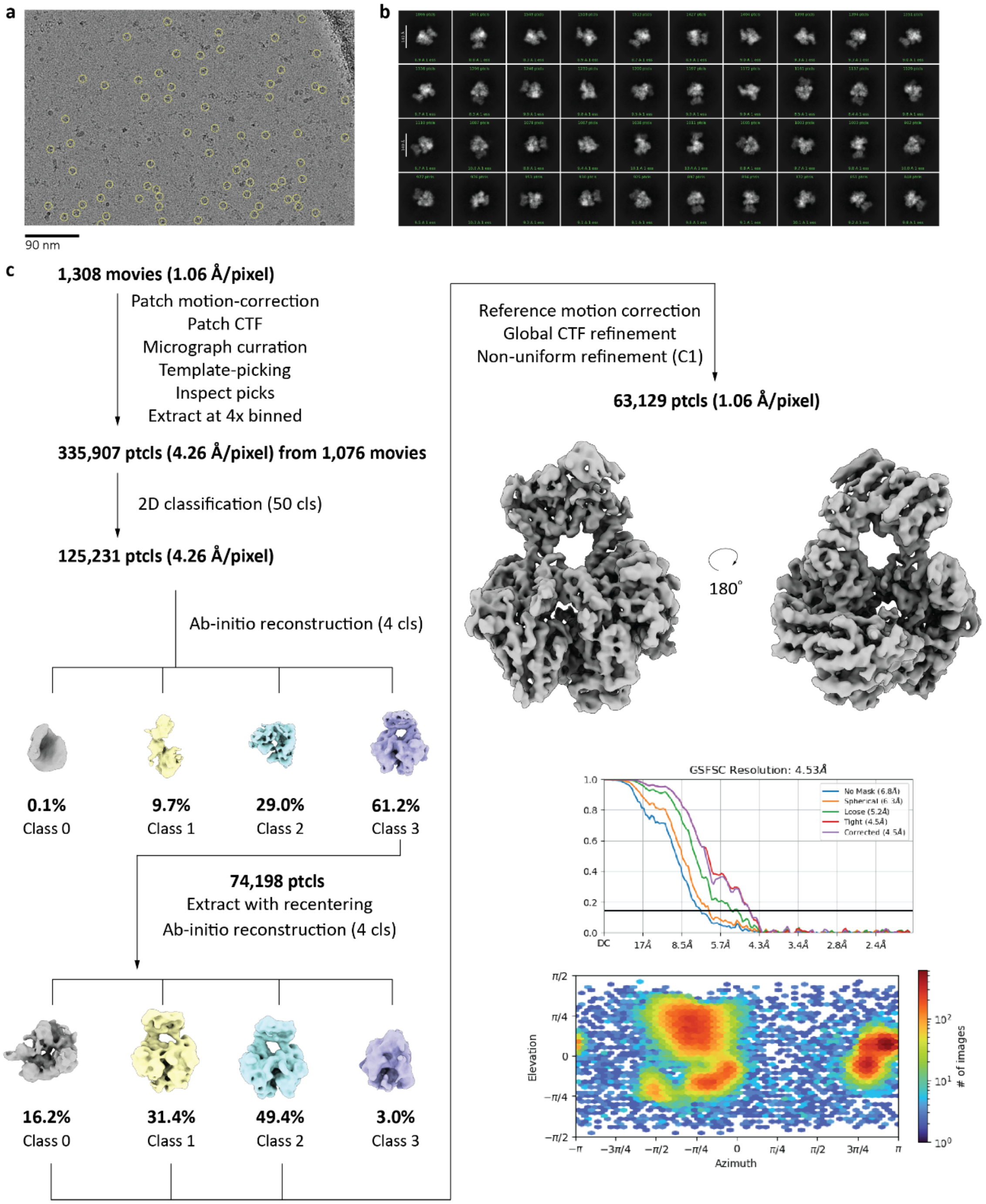

**Extended Data Figure 6 – Data processing pipeline for PP and RvLEAM$_{short}$–(1:6) dataset**
1 µM PP with 6 µM RvLEAM$_{short}$ **(a)** Representative micrograph. **(b)** Top 40 classes from 2D classification of PP. **(c)** Cryo-EM processing pipeline used to obtain final cryo-EM map for PP.

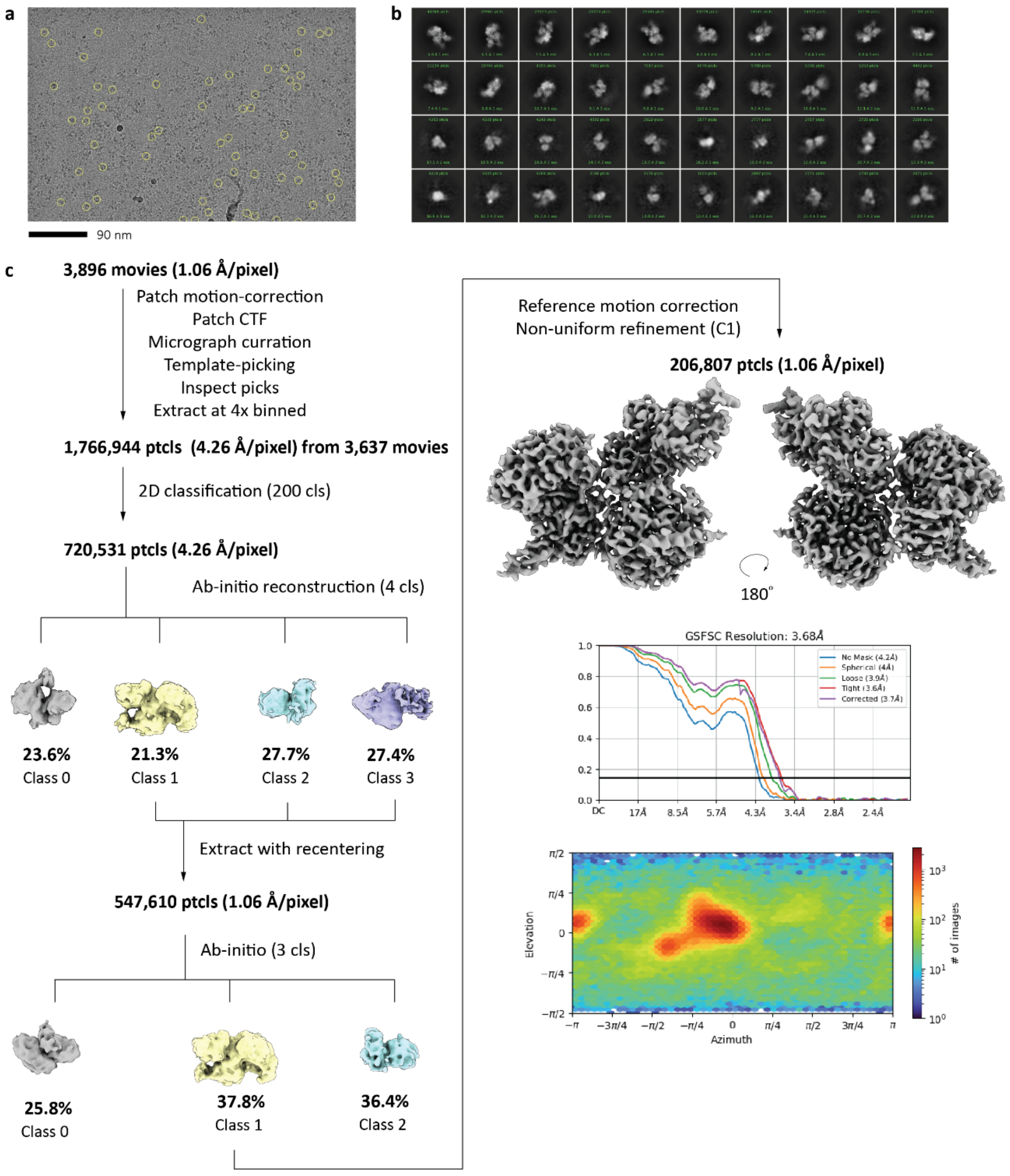

**Extended Data Figure 7 – Data processing pipeline for PRC2 and RvLEAM$_{\mathrm{short}}$ (1:6) dataset** 1 µM PRC2 with 6 µM RvLEAM$_{\mathrm{short}}$ **(a)** Representative micrograph. **(b)** Top 40 classes from 2D classification of PRC2. **(c)** Cryo-EM processing pipeline used to obtain final cryo-EM map of PRC2.

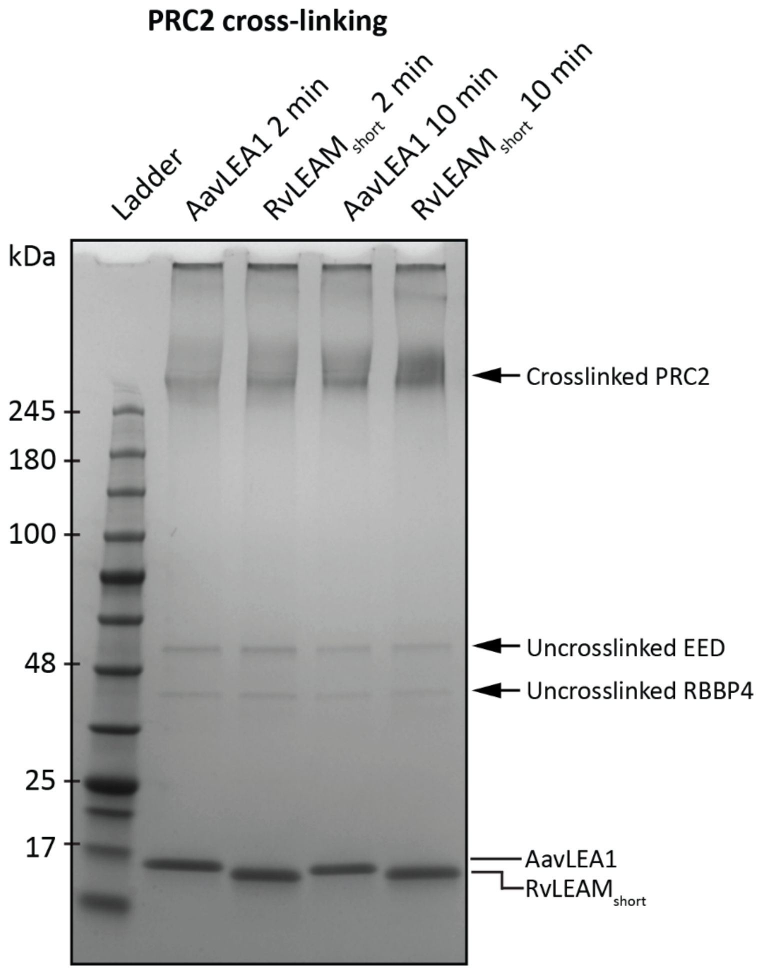

**Extended Data Figure 8 – PRC2 cross-linking validation SDS-PAGE** SDS-PAGE analysis of cross-linked PRC2 with AavLEA1 (1:12) or RvLEAM (1:6) after 2 and 10 minutes incubation with glutaraldehyde. Labels denoted with a * are predicted subunits of PRC2 based on molecular weight.

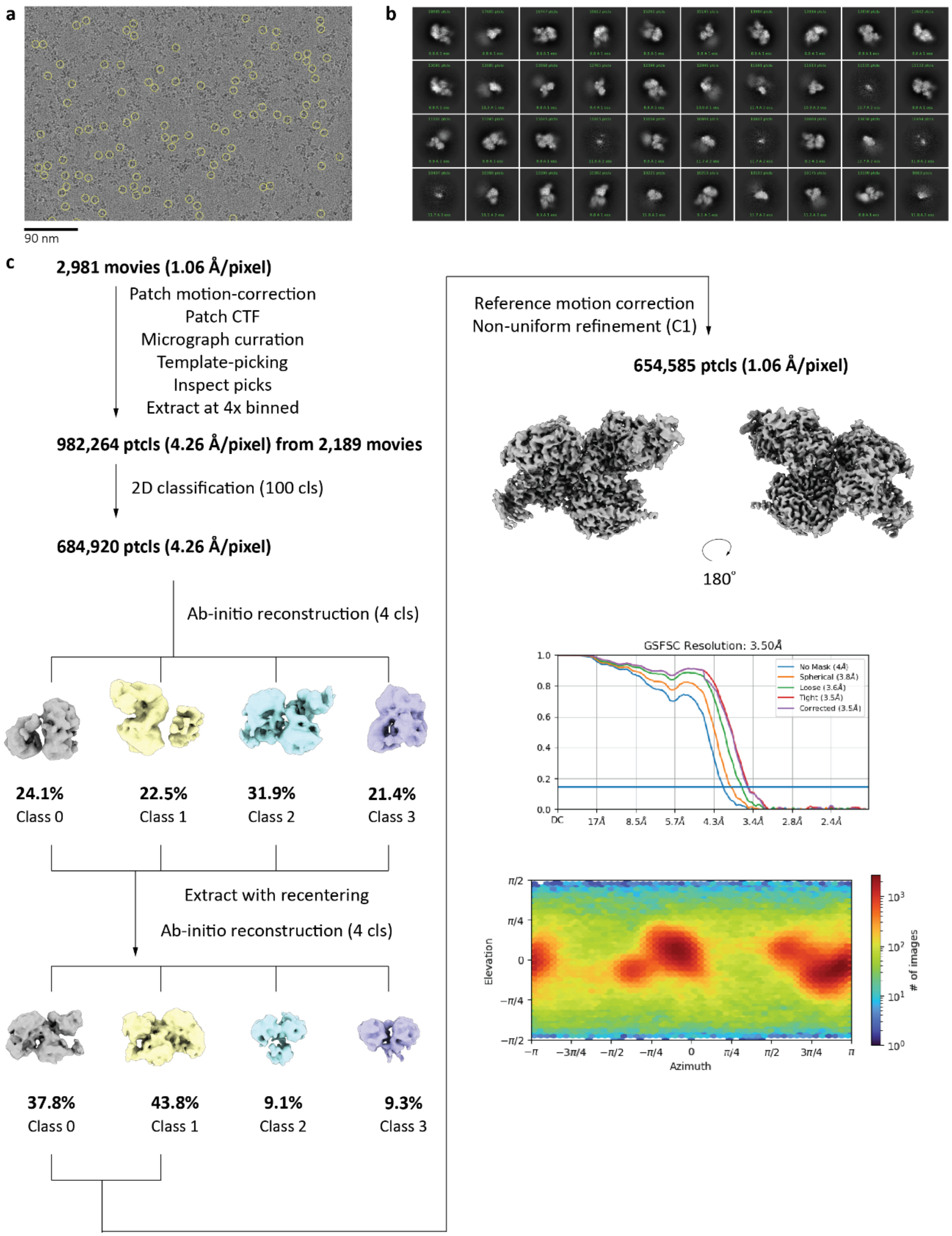

**Extended Data Figure 9 – Data processing pipeline for PRC2 and RvLEAM$_{short}$ (1:6) dataset with mild cross-linking (2 minutes)** 1 µM PRC2 with 6 µM RvLEAM$_{short}$ **(a)** Representative micrograph. **(b)** Top 40 classes from 2D classification of PRC2. **(c)** Cryo-EM processing pipeline used to obtain final cryo-EM map of PRC2.

**Table 1: Cryo-EM data collection, refinement, and validation statistics**

| | PP, AavLEA1 (1:40) (EMD-43628) (PDB 8VY3) |
|---|---|
| **Data collection and processing** | |
| Magnification | 79,000 |
| Voltage (kV) | 200 |
| Electron exposure (e–/Å$^2$) | 50 |
| Defocus range (µm) | -2.5 – (-1.0) |
| Pixel size (Å) | 1.064 |
| Symmetry imposed | C1 |
| Initial particle images (no.) | 2,930,318 |
| Final particle images (no.) | 856,205 |
| Map resolution (Å) | 2.98 |
| FSC threshold | 0.143 |
| Map resolution range (Å) | |
| **Refinement** | |
| Initial model used (PDB code) | 5EXR |
| Model resolution (Å) | 3.0 |
| FSC threshold | 0.143 |
| Model resolution range (Å) | |
| Map sharpening *B* factor (Å$^2$) | |
| Model composition | |
| Non-hydrogen atoms | 18829 |
| Protein residues | 2324 |
| Ligands | SF4:1, Zn:3 |
| *B* factors (Å$^2$) | |
| Protein | 53.58 |
| Ligand | 56.15 |
| R.m.s. deviations | |
| Bond lengths (Å) | 0.004 (0) |
| Bond angles (°) | 0.760 (2) |
| Validation | |
| MolProbity score | 1.79 |
| Clashscore | 6.26 |
| Poor rotamers (%) | 1.61 |
| Ramachandran plot | |
| Favored (%) | 95.83 |
| Allowed (%) | 3.95 |
| Disallowed (%) | 0.22 |
